# Thiolation-Based Protein-Protein Hydrogels for Improved Wound Healing

**DOI:** 10.1101/2023.10.19.563054

**Authors:** Xing Liu, Jie Wang, Zhao Guo, Wenting Shen, Zhenzhen Jia, Shuang Jia, Limiao Li, Jieqi Wang, Liping Wang, Jiaqi Li, Yufang Chen, Yinan Sun, Min Zhang, Jia Bai, Liyao Wang, Xinyu Li

## Abstract

The limitations of protein-based hydrogels, including their insufficient mechanical properties and restricted biological functions, arise from the highly specific functions of proteins as natural building blocks. A potential solution to overcome these shortcomings is the development of protein-protein hydrogels, which integrate structural and functional proteins. In this study, we introduce a protein-protein hydrogel formed by crosslinking bovine serum albumin (BSA) and a genetically engineered intrinsically disordered collagen-like protein (CLP) through Ag-S bonding. Our approach involves thiolating lysine residues of BSA and crosslinking CLP with Ag^+^ ions, utilizing thiolation of BSA and the free-cysteines of CLP. The resulting protein-protein hydrogels exhibit exceptional properties, including notable plasticity, inherent self-healing capabilities, and gel-sol transition in response to redox conditions. Furthermore, in comparison to standalone BSA hydrogels, these protein-protein hydrogels demonstrate remarkable antibacterial properties, enhanced cellular viability, and improved cellular migration. In vivo experiments provide conclusive evidence of accelerated wound healing, observed not only in murine models with streptozotocin (Step)-induced diabetes but also in zebrafish models subjected to UV-burn injuries. Detailed mechanistic insights, combined with assessments of pro-inflammatory cytokines and the expression of epidermal differentiation-related proteins, robustly validate the protein-protein hydrogel’s effectiveness in promoting wound repair. This pioneering approach advances the development of protein-protein hydrogels and serves as a reference for the creation of multifunctional protein-based hydrogels.

## 1. Introduction

Proteins, as fundamental building blocks of life, have garnered increasing attention in the development of bioactive materials[1-5]. Protein-based biomaterials, known for their high biocompatibility and direct therapeutic effects, have emerged as promising candidates for a wide spectrum of applications in regenerative medicine[6, 7], disease treatment[8, 9], and tissue engineering[2, 5, 10-12]. Protein-based hydrogels, in particular, have demonstrated remarkable potential in various tissue engineering endeavors, such as wound healing[2, 13, 14] and cartilage regeneration[3, 15].

Due to their exceptional biological specificity, proteins that exhibit both excellent hydrogel scaffolding properties and highly specific biological functions have not yet been reported. For example, fibroblast growth factor (bFGF) is excellent at promoting wound healing but cannot function as a structural protein for hydrogel preparation[16]. Bovine serum albumin (BSA) is commonly used as a structural protein for protein hydrogel preparation; however, its lack of biological activity limits its further applications[17, 18]. Therefore, the development and application of protein-based hydrogels can greatly benefit from the crosslinking of two or more proteins, simultaneously introducing a high physical property scaffold and bioactive functional groups[4, 19].

The choice of a crosslinking strategy plays a pivotal role in achieving protein-protein crosslinking and hydrogel preparation[2, 3, 20, 21]. While various techniques have been developed for protein crosslinking or assembly[4, 10], some strategies like unfold-chemical coupling crosslinking are not suitable for protein-protein hydrogels preparation due to their harsh reaction conditions[22], which can disrupt the protein structure and lead to a loss of function.

Among amino acid-based conjugation strategies[20, 23], cysteine has been commonly targeted for crosslinking, often involving Thiol-Michael addition reactions[2, 19, 20, 23, 24]. The high nucleophilicity and responsiveness to metal ions makes cysteine an attractive option for crosslinking[14, 17]. However, the limited presence of native thiols in many proteins and the potential disruption of protein structure due to disulfide bond reduction have hindered the use of cysteine-based crosslinking in recent hydrogel development[2, 20, 24]. An alternative modification and crosslinking site on proteins is lysine, characterized by its ε-amine[2, 20, 25]. Lysine is frequently targeted using activated esters, such as N-hydroxysuccinimide (NHS) [20], in combination with multifunctional cross-linkers like β-[tris(hydroxylmethyl) phosphino] propionic acid (THPP) [26] and tetrakis (hydroxymethyl) phosphonium chloride (THPC) [27]. The nucleophilicity of the ε-amine, located on its side chain, along with its ubiquity in the human proteome (approximately 6% of residues), makes lysine an attractive option for crosslinking[25]. However, crosslinking or modification of the primary amines on lysine residues may impact protein function, as lysine often forms part of the catalytic triad in enzymes[20].

Above all, the development of protein-protein hydrogels has posed a significant challenge, limiting the progress in the field of protein-based hydrogels. To overcome this obstacle, we employed Traut’s reagent to thiolate lysine residues in BSA. Traut’s reagent is well-known for its capacity to add thiol groups to the primary amines (-NH2) of lysine, thereby preserving charge properties akin to the original amino group[14, 17, 28, 29]. The thiolated BSA, functioning as a globular protein scaffold, was crosslinked with a genetically engineered collagen-like protein (CLP). The CLP was deliberately designed to include free cysteines and enhance wound healing[24]. Through the addition of Ag^+^, the thiolated BSA and cysteine-free CLP formed a hydrogel via Ag-S bonding. This strategy not only enabled the preparation of protein-protein hydrogels with exceptional mechanical properties but also harnessed the combined advantages of BSA, CLP and Ag^+^. We conducted extensive investigations at both cellular and in vivo levels to evaluate their therapeutic potential. In summary, our study presents a novel strategy for the preparation of protein-protein hydrogels and introduces a protein-based hydrogel with promising applications in wound healing.

## 2. Results

### 2.1 Hydrogel preparation

To create protein-protein hydrogels, we opted for bovine serum albumin (BSA) as the structural protein due to its documented exceptional mechanical properties[22, 30]. For the biofunctional component, we genetically engineered a cysteine-free collagen-like protein (CLP)[24] **(Fig 1 A and Supplementary Fig 1)**. The CLP features two free cysteine residues at the N- and C-terminals **(Fig 1 A D)**. Additionally, to enhance the stability of the triple-helix structure of CLP, two CPPC peptides were incorporated at the N- and C-terminals of the collagen-like domains **(Fig 1 A)**. We integrated RGD sequences into the protein to facilitate cell adhesion. The identity of the CLP was further verified through SDS-page **(Fig 1 B)** and Western Blot **(Fig 1 C)** using an anti-Histag antibody. Its molecular weight, as determined by MALDI-TOF, was measured at 36.38 kDa **(Supplementary Fig 3)**.

**Figure 1.**
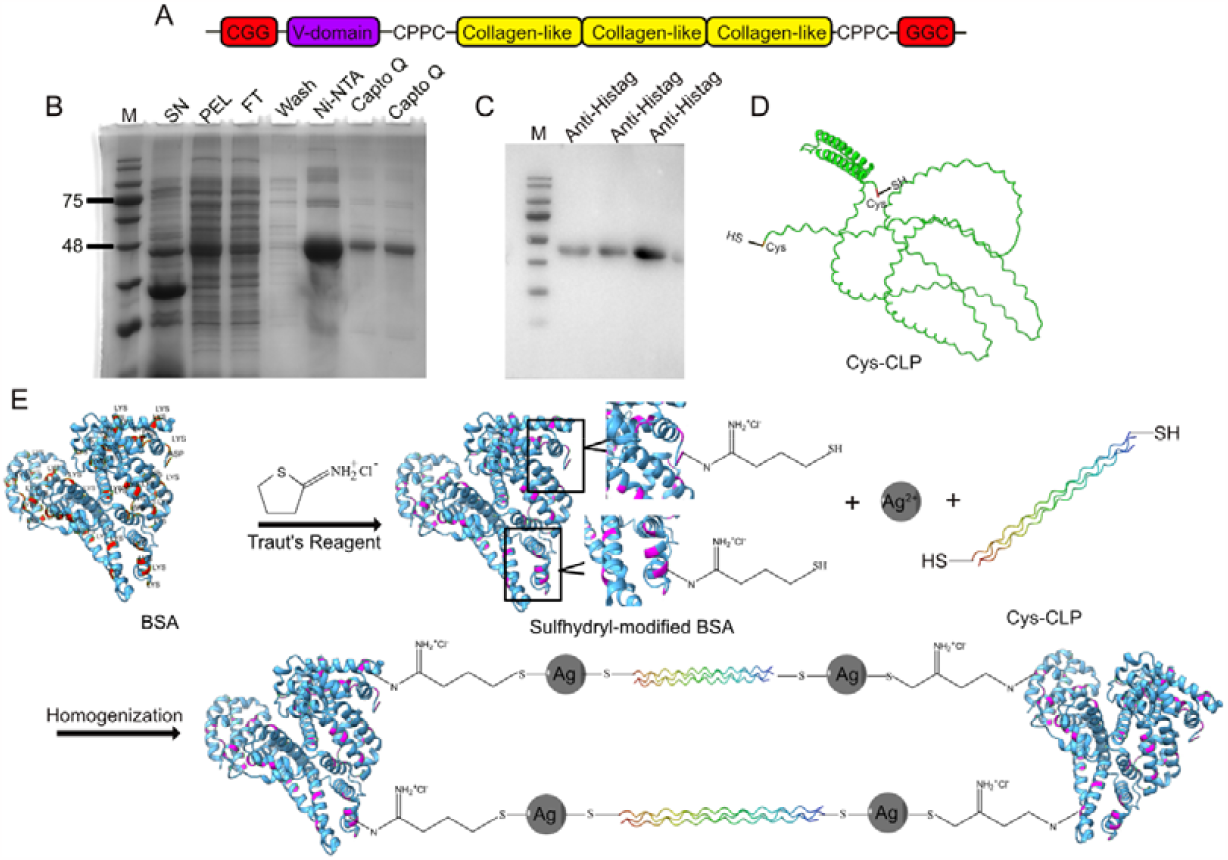
CLP construction and mechanisms of hydrogel formation. A. Genetic blueprint for CLP. B. SDS-PAGE analysis demonstrating successful CLP purification. C. Western blot to verify the integrity of the purified CLP. D. Structural representation of CLP modeled using Alphafold. B. Mechanisms of crosslinking BSA thiol groups and BSA-Ag-CLP hydrogel formation.

Since the BSA has limited free cysteine residues due to their role in forming disulfide bonds for structural stability[31], we utilized Traut’s reagent to introduce thiol groups to the BSA. Following modification and lyophilization, the thiolated BSA was reconstituted in a sodium acetate/acetic acid solution, and Ag^+^ was added to create the BSA hydrogel (BSA-Ag-BSA) as the control group for the protein hydrogel containing only the structural protein **(Supplementary Fig 2)**. In the case of the protein-protein hydrogel, the CLP solution was combined with the thiolated BSA solution in the same buffer. Subsequently, Ag^+^ was introduced into the mixture, leading to the immediate formation of the protein-protein hydrogel (BSA-Ag-CLP) **(Fig 1 E)**.

### 2.2 Apparent properties of prepared hydrogels

In accordance with this strategy, we have successfully synthesized two distinct hydrogels, denoted as BSA-Ag-BSA and BSA-Ag-CLP **(Fig 2 A)**. The crosslinking process, relying on the presence of free cysteines and lysine modification, had minimal impact on the bioactivities of the two proteins. Upon the introduction of Ag^+^ ions, the thiol-modified protein solution underwent a rapid transformation into structurally sound protein hydrogels **(Fig 2 B)**. Notably, an in-depth analysis employing scanning electron microscopy (SEM) unveiled a consistent and well-defined pore network microstructure within these hydrogels **(Fig 2C)**. This result indicates that BSA, serving as the structural protein, contributes excellent hydrogel structural characteristics, while the introduction of CLP does not induce significant alterations in the hydrogel structure. Furthermore, these hydrogels demonstrated noteworthy non-Newtonian fluid characteristics, thus corroborating their suitability for injectability applications **(Fig 2D)**. They exhibited robust tissue adhesion properties, securely maintaining their attachment even under rigorous testing conditions, such as extrusion, vertical compression, and twisting on cowhide surfaces **(Fig 2E)**. Intriguingly, we observed that the optical transmittance of these hydrogels exhibited a pH-dependent behavior, showing a discernible decrease as the pH levels increased **(Fig 2F)**. These highly flexible hydrogels are suitable for diverse wound shapes and can be molded accurately and removed without distortion in hours **(Fig 2G)**. These hydrogels demonstrated an inherent ability to self-heal, most likely due to the dynamic formation of Ag-S coordination bonds, which imparts excellent self-healing properties[32] **(Fig 2H)**. The Ag-S bonds are sensitive to redox reactions, leading us to explore their responsiveness. Both protein hydrogels exhibited gel-sol transitions upon exposure to DTT and H_2_O_2_ **(Fig 2I)**, with BSA-Ag-CLP showing a faster transition due to CLP’s flexible structure.

**Figure 2.**
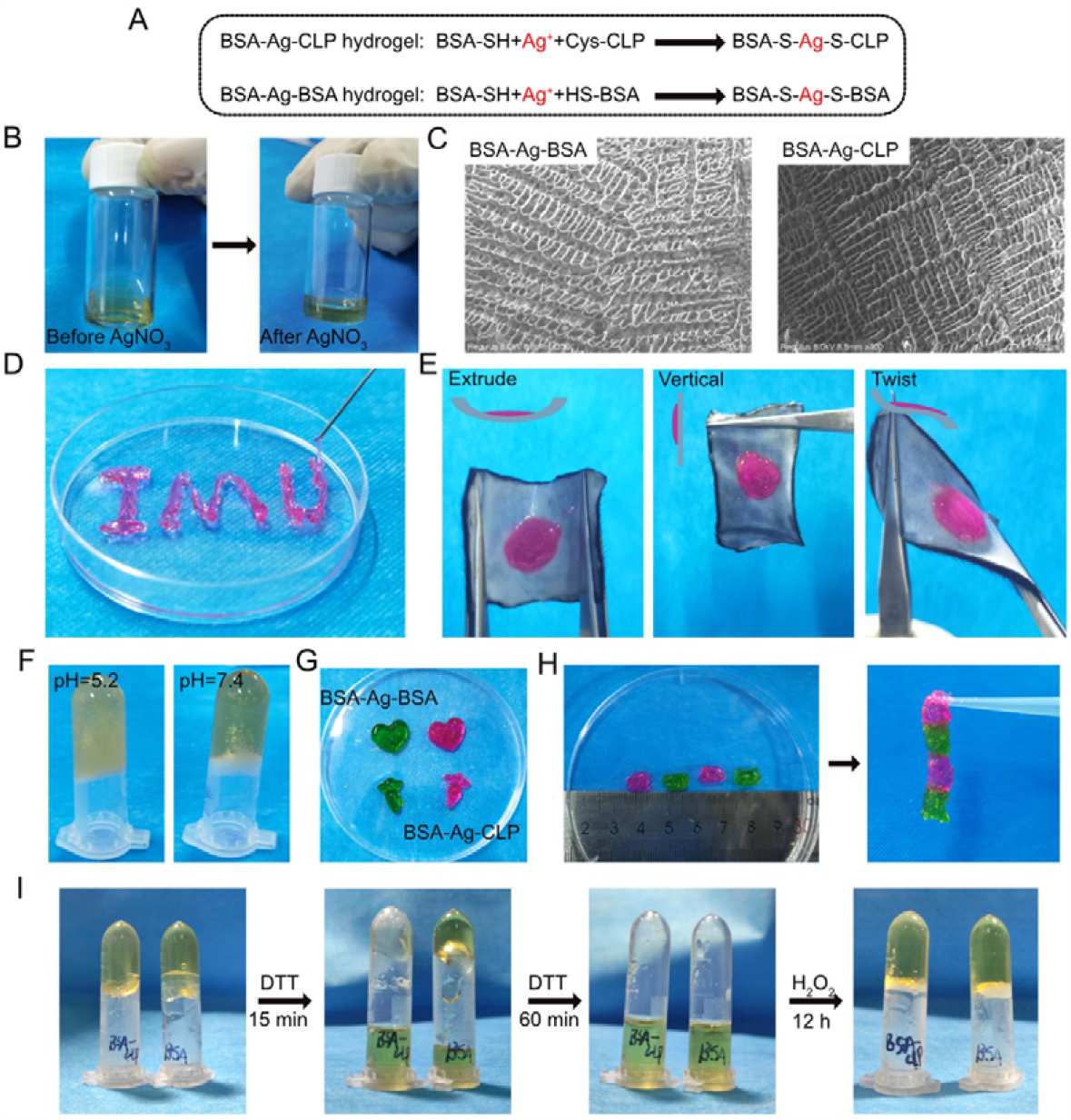
Apparent Properties of Prepared Hydrogels. A. Protein-protein hydrogels synthesized in this study. B. Photographic representation of pre-hydrogel and formed hydrogels. C. images illustrating the microstructure of BSA-Ag-BSA and BSA-Ag-CLP hydrogels. D. Demonstrated injectability of the hydrogel. E. Examination of hydrogel adhesion on cowhide. F. Evaluation of hydrogel performance under different pH conditions. G. Assessment of hydrogel plasticity. H. Investigation of the self-healing capability of the hydrogel. I. Oxidation-reduction responsive properties of prepared hydrogels.

### 2.3 Characterization of Prepared Hydrogels

The characteristics of the protein hydrogels were investigated using FTIR, rheological analysis, and circular dichroism (CD). FTIR analysis showed peaks at 1644 cm^-1^ and 1390 cm^-1^, indicating structural changes in the formation of BSA-Ag-CLP hydrogel. The appearance of the 1644 cm^-1^ peak suggested increased α-helix content **(Fig 3A)**, while changes at 1390 cm^-1^ indicated alterations in amino acid side-chain conformations **(Fig 3A)**. These findings highlight the enhanced stability of CLP through its crosslinking with BSA, considering that CLP is intrinsically disordered. The increased peak intensity at 220 nm in the CD results confirms the improved stability of CLP, as it indicates alterations in the secondary structure introduced by CLP when compared to the BSA-Ag-BSA hydrogel **(Fig 3D)**. Rheological analysis showed that BSA-Ag-BSA hydrogel exhibited higher mechanical properties compared to BSA-Ag-CLP, likely due to the incorporation of intrinsically disordered proteins (CLP), enhancing hydrogel flexibility **(Fig 3B C)**.

**Figure 3.**
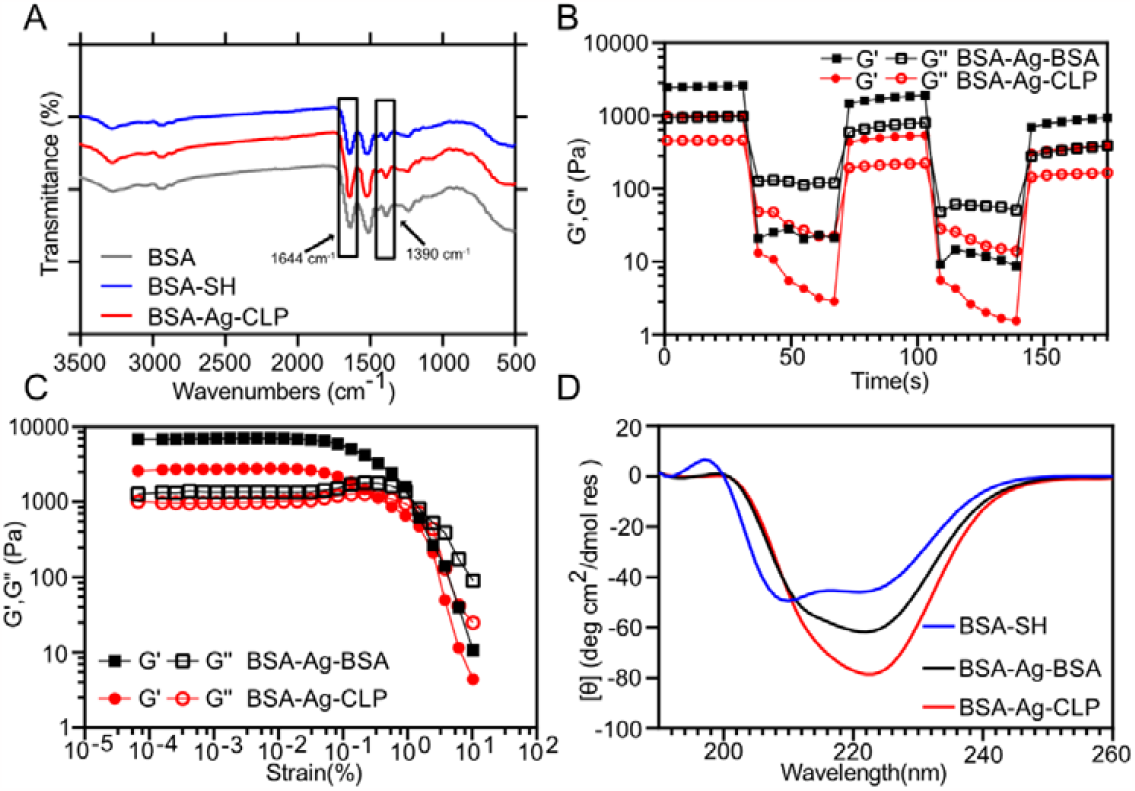
Characterization of Protein Hydrogels. A. FTIR analysis of hydrogels. B. Dynamic step-strain measurements of the hydrogels. C. Strain Sweep of hydrogels. D. CD analysis revealing changes in protein folding.

### 2.4 Antibacterial Properties of Protein Hydrogels

The presence of Ag suggested potential antibacterial properties in the resulting protein hydrogels[17]. To assess these properties, we conducted antibacterial assays using agar plates and optical density (OD) measurements. Our results demonstrated a significant antibacterial effect on agar plates, affecting both the Gram-negative bacterium *Escherichia coli* and the Gram-positive bacterium *Staphylococcus aureus* **(Fig 4A)**. Furthermore, we examined the antibacterial activity of the hydrogels under culture conditions, measuring OD at two-hour intervals. In comparison to the control group, the inhibition rate rapidly increased within the first four hours, reaching 80% for *E. coli* and 90% for *S. aureus* **(Fig 4B)**. Interestingly, no significant differences were observed between the BSA-Ag-BSA and BSA-Ag-CLP hydrogels, due to their equivalent Ag^+^ concentrations **(Fig 4B)**.

**Figure 4.**
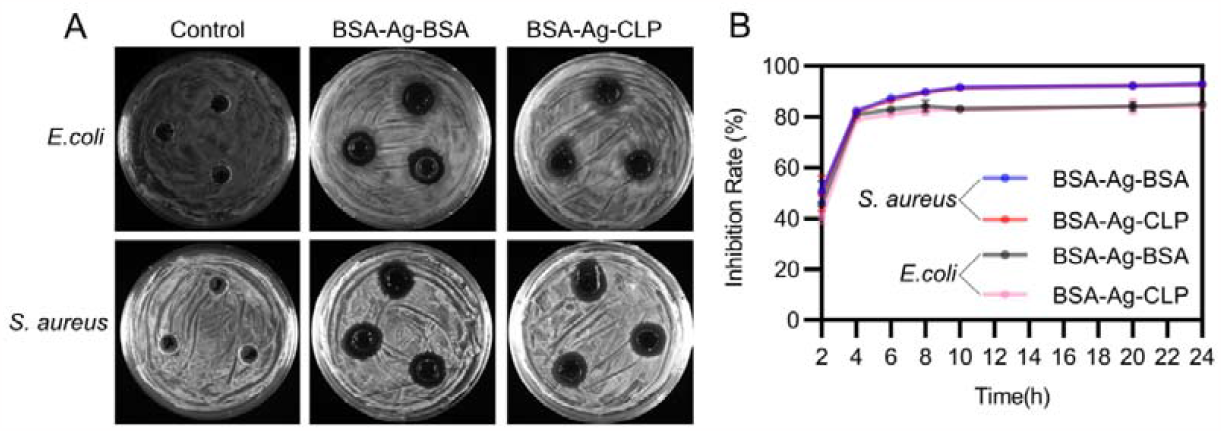
Antibacterial Effects of Protein Hydrogels. A. Antibacterial assays on agar plates. B. Antibacterial activity under culture conditions, as measured by optical density (OD) at two-hour intervals. All experiments with three repeats.

### 2.5 Biocompatibility evaluation of hydrogels in vitro

In vitro biocompatibility assessment of the two prepared hydrogels was conducted using NIH-3T3 and HaCat cell lines. Cell viability was determined through the Cell Counting Kit-8 assay post-hydrogel treatment. Cells were co-incubated with BSA-Ag-BSA and BSA-Ag-CLP at concentrations of 1%, 3%, and 5% (v/v), respectively, with PBS at matching concentrations serving as the control group. The results revealed a significant increase in cell viability for both cell lines following hydrogel treatment **(Fig 5A)**. Notably, BSA-Ag-CLP exhibited a more pronounced enhancement of cell viability compared to BSA-Ag-BSA, suggesting that the introduction of CLP may have a less potent effect on promoting cell viability than BSA alone **(Fig 5A)**. To further visualize cell viability, we performed live/dead staining experiments using 5% hydrogel treatments. The results indicated minimal dead cells and an increase in live cell density for both NIH-3T3 and HaCat cells after hydrogel treatment **(Fig 5B)**. These findings align with similar trends observed in cell scratch assays, where the BSA-Ag-CLP-treated cells exhibited a higher rate of wound healing compared to the BSA-Ag-BSA-treated group **(Fig 5C)**.

**Figure 5.**
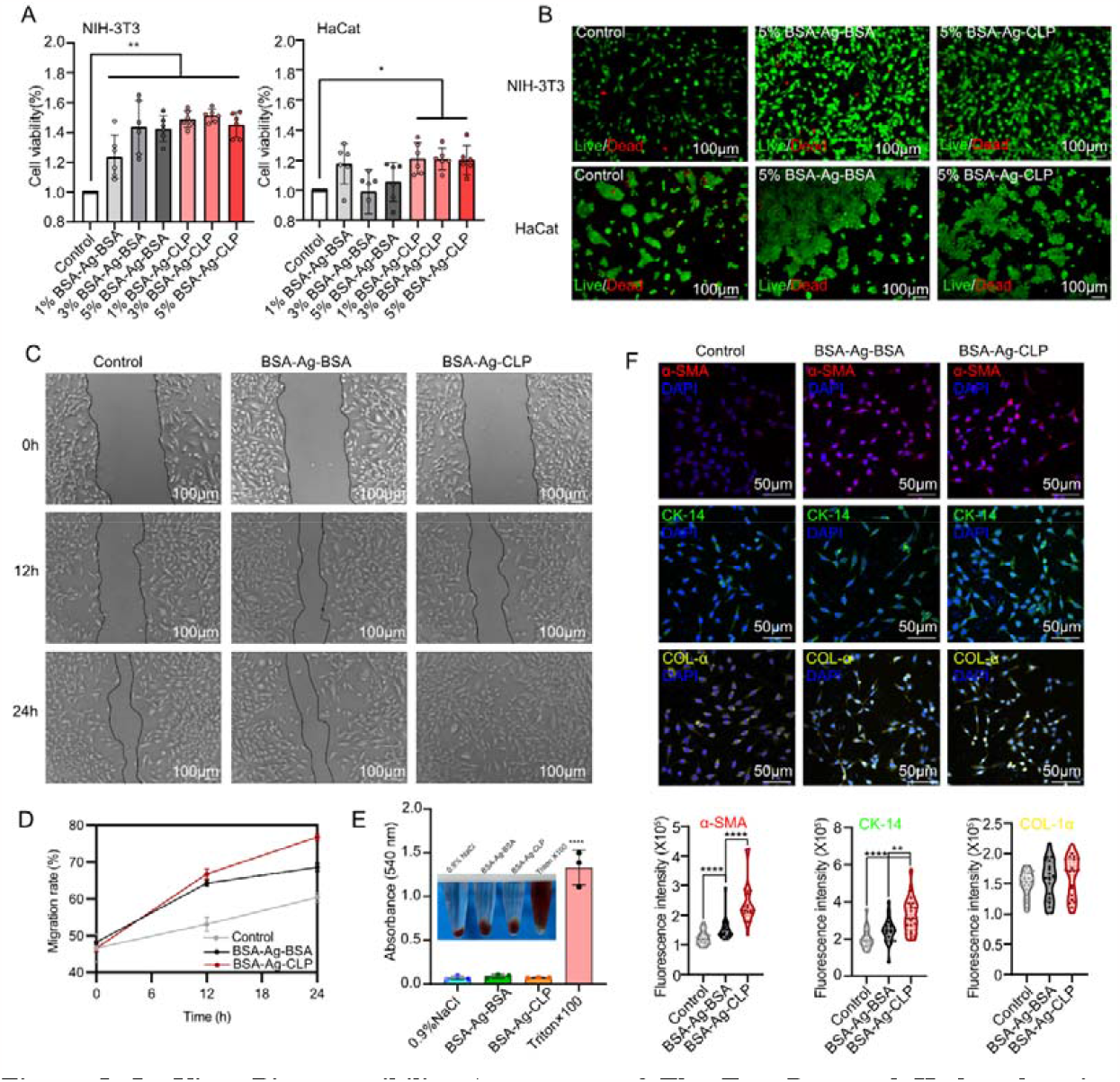
In Vitro Biocompatibility Assessment of The Two Prepared Hydrogels using NIH-3T3 and HaCat Cell Lines. A. Cell viability of NIH-3T3 and HaCat cell assessed by the CCK-8 assay. B. Live/dead staining experiments of NIH-3T3 and HaCat cells. C. Cell scratch images and quantitative analysis of cell wound healing rate. D. Immunofluorescence assays for α-SMA, CK-14, and COL-1α. N=6, Data are presented as means ± SD. ^*^p < 0.05, ^**^p < 0.01, ^***^p < 0.001.

α-Smooth muscle actin protein (α-SMA) serves as a well-established marker for smooth muscle cells, particularly in studies related to tissue regeneration[33-35]. In the context of epidermal regeneration, collagen-1α (COL-1α) and cytokeratin 14 (CK-14) collaborate to maintain skin structure and function, thus promoting wound healing and the regeneration of epidermal cells[14, 35]. Immunofluorescence assays were therefore performed to assess the protein-level effects of the hydrogels on NIH-3T3 cells, specifically targeting α-SMA, CK-14, and COL-1α. The findings revealed that the administration of both BSA-Ag-BSA and BSA-Ag-CLP hydrogels led to an increase in the expression of α-SMA, CK-14, and COL-1α **(Fig 5F)**. These findings underscore the enhanced protein expression in NIH-3T3 cells due to the introduction of CLP, which has the potential to positively impact epidermal cell regeneration promoted by the protein hydrogels.

Furthermore, we conducted investigations into the hemolysis and hemostatic properties of the hydrogels[36, 37]. The results indicated that when the hydrogels were incubated with fresh erythrocyte suspension, the clarity of the supernatant resembled that of the 0.9% saline group **(Fig 5E)**. This suggests that the hydrogel exhibits favorable cytocompatibility and blood compatibility as a wound dressing. Moreover, we assessed the rapid hemostatic properties of the hydrogels using a mouse liver bleeding model, where the hydrogels significantly reduced blood loss **(Supplementary Fig 4)**.

### 2.6 Enhanced Wound Healing in UV-Burned Zebrafish Model

To assess the wound healing efficacy of the hydrogels, we established a UV-burned zebrafish model[38] and applied the hydrogels to the burn site every three days. The results revealed that in the BSA-Ag-CLP group, the UV-burned wounds in zebrafish had nearly completely healed after 18 days, with significantly smaller wounds compared to the BSA-Ag-BSA group **(Fig 6A)**. This enhanced wound healing can be attributed to the introduction of CLP. Additionally, both hydrogel treatment groups exhibited higher wound healing rates than the control group, demonstrating the wound-recovery-promoting effect of hydrogel application **(Fig 6A)**. After the 18-day treatment period, we euthanized the zebrafish for pathological analysis. We conducted Hematoxylin and Eosin (H&E) staining and Masson’s trichrome staining to assess tissue regeneration and collagen deposition **(Fig 6B)**. Consistent with the visual observations, the BSA-Ag-CLP treatment exhibited less collagen deposition and the highest recovery rate compared to the other groups **(Fig 6B)**.

**Figure 6.**
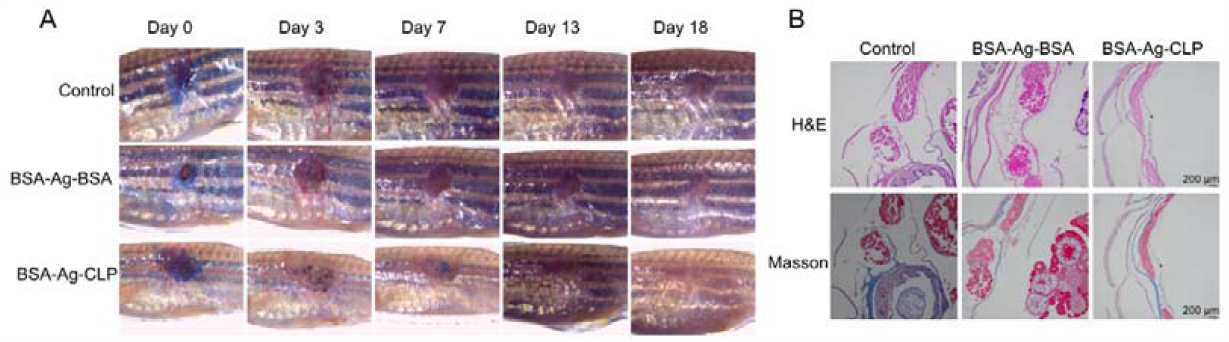
Evaluation of Hydrogel-Mediated Wound Healing in a UV-Burned Zebrafish Model. A. Microscopic images illustrating the wound healing process in zebrafish treated with BSA-Ag-CLP and BSA-Ag-BSA hydrogels. B. Histological evaluation of tissue regeneration and collagen deposition, conducted through H&E staining and Masson’s trichrome staining. N=7.

### 2.7 Enhanced wound healing in diabetic mouse model

We assessed the wound healing efficacy of the hydrogels in a diabetic mouse model[24]. The mice were initially administered streptozotocin (Step) for 7 days to induce hyperglycemia, and blood glucose levels were monitored (**Fig 7A)**. After 7 days of injections, the STZ-injected group exhibited an average blood glucose level of approximately 15 mmol/L, significantly higher than the saline-injected control group (5 mmol/L) (**Fig 7B)**, confirming the successful establishment of the diabetic mouse model.

**Figure 7.**
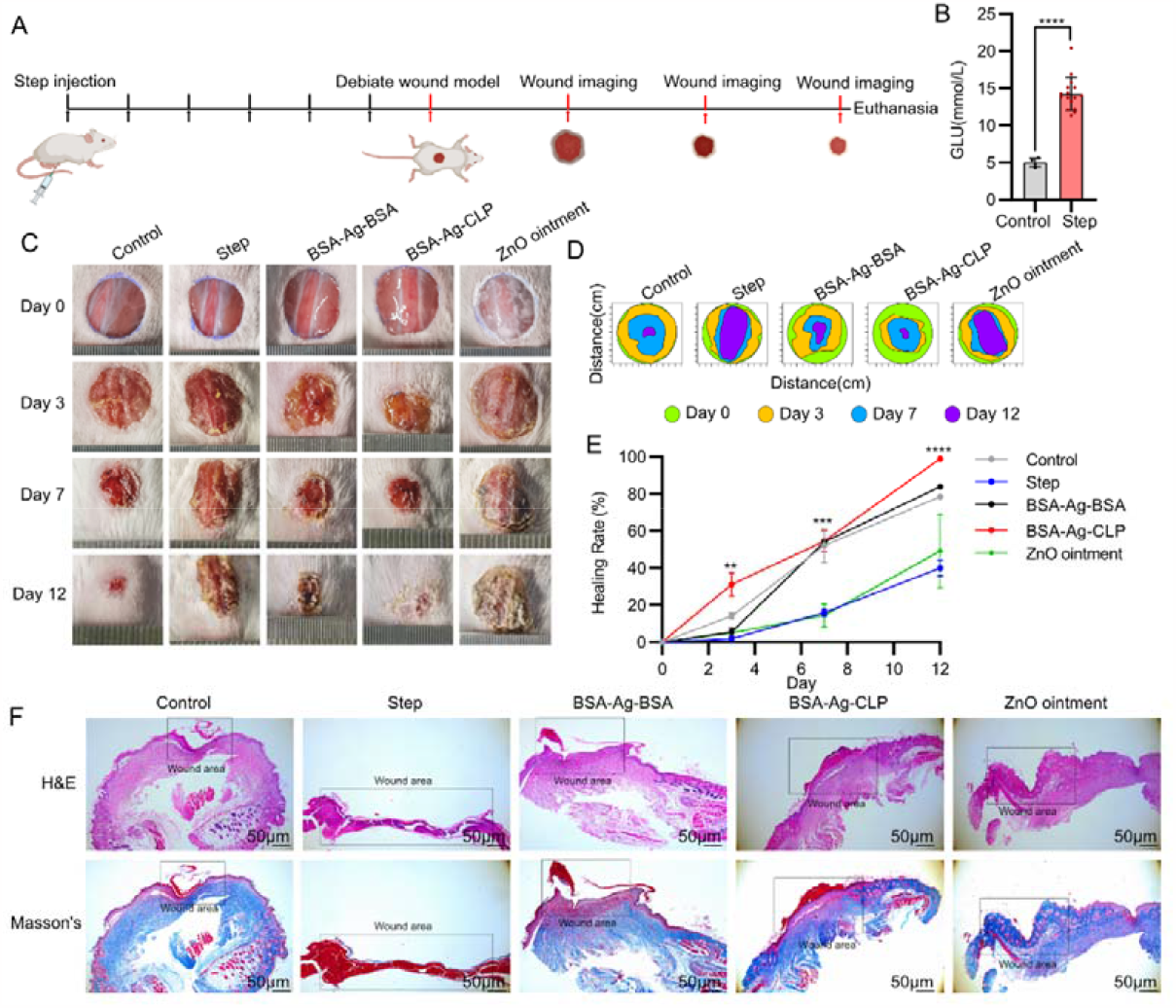
Assessing Hydrogel-Mediated Wound Healing in a Diabetic Mouse Model. A. Experimental design and workflow. B. Blood glucose levels at day 7 post-Step injection. C. Representative images of wound healing. D. Comparison of wound healing boundaries among different groups. E. Wound healing rates at various time points across different groups. F. Histological evaluation of wound pathology using H&E staining and Masson’s trichrome staining. N=4, data are presented as means ± SD. ^*^p < 0.05, ^**^p < 0.01, ^***^p < 0.001.

Subsequently, the mice were divided into five groups, and 1 cm diameter wounds were created on their dorsal surfaces to simulate diabetic wounds. Wound progress was monitored on days 0, 3, 7, and 12 post-wounding (**Fig 7A)**. The results indicated a significant improvement in wound healing following hydrogel administration. In the BSA-Ag-CLP group, diabetic wounds nearly completely closed within 12 days (**Fig 7B C and E)**. Although the wound healing rate in the BSA-Ag-BSA groups was lower than in the BSA-Ag-CLP group, it was significantly higher than in the Step-induced diabetic group (**Fig 7B C and E)**. Interestingly, both hydrogel-treated groups exhibited better wound healing than the positive control group treated with ZnO ointment (**Fig 7B C and E)**.

Furthermore, pathological analysis involving H&E staining and Masson’s trichrome staining was conducted to assess wound pathology (**Fig 7F)**. Both hydrogel-treated groups showed minimal scarring and skin recovery beneath the scar, while the Step group exhibited persistent scarring without skin recovery (**Fig 7F)**. In contrast, the ZnO ointment group displayed no scar formation, resulting in an unrecovered wound (**Fig 7F)**. These results further support the notion that hydrogel application improves wound healing in the diabetic wound model.

### 2.8 Immunofluorescence and Western Blot Analysis

To understand the mechanisms underlying hydrogel-facilitated wound healing, we first assessed the levels of α-SMA, COL-1α, and CK-14 at the wound site through immunofluorescence. The results revealed a significant increase in the expression of these three proteins in the hydrogel-treated groups compared to the Step-induced diabetic and ZnO ointment groups (**Fig 8A)**. This was further validated by Western blot analysis of the entire wound site (**Fig 8B)**, confirming that hydrogel application leads to changes in protein expression, ultimately enhancing wound healing. Importantly, the BSA-Ag-CLP group exhibited notably higher protein expression than the BSA-Ag-BSA group, underscoring the beneficial impact of CLP as a functional substrate in improving protein expression.

**Figure 8.**
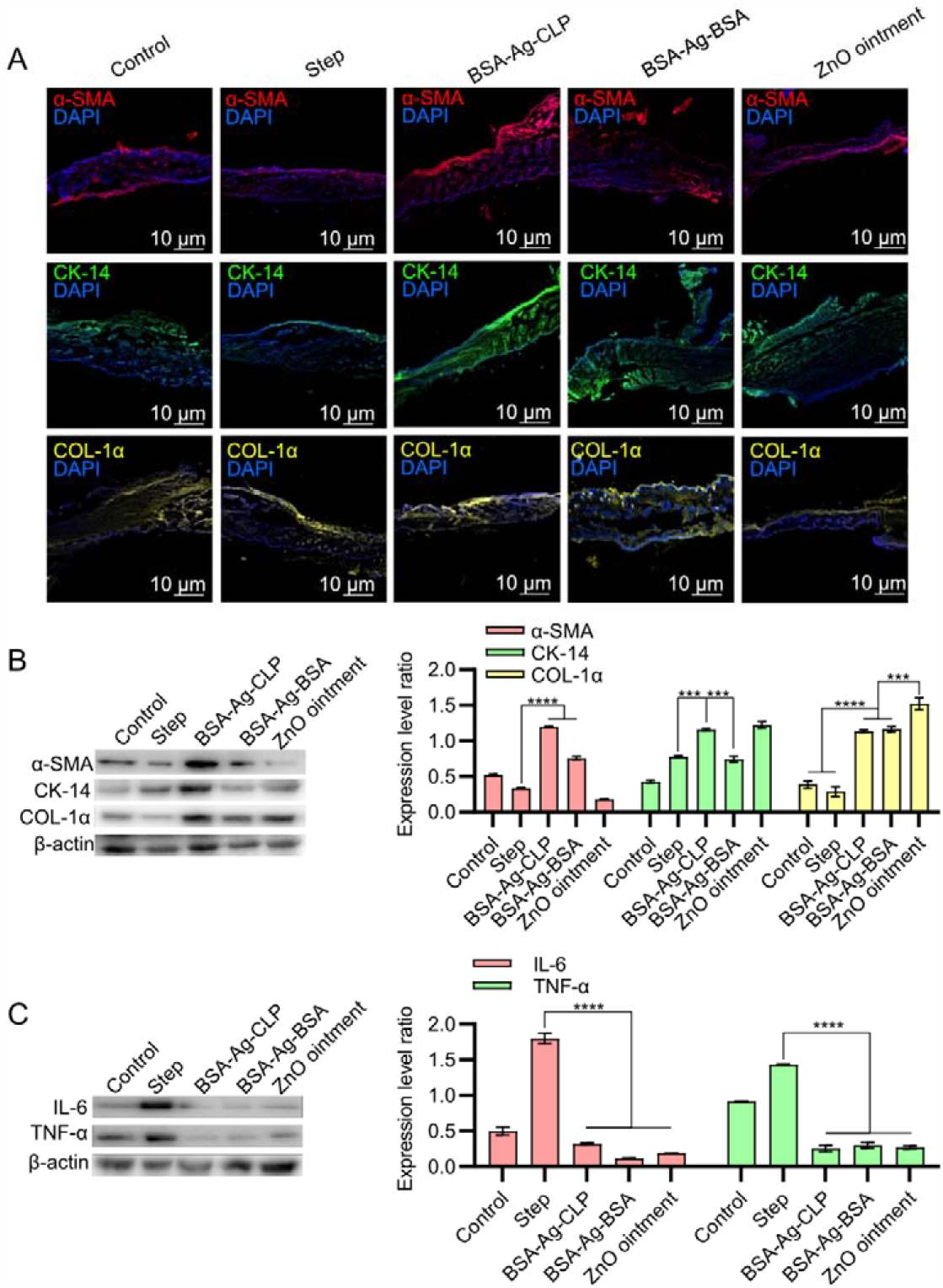
Mechanisms Underlying Hydrogel-Mediated Wound Healing. A. Immunofluorescence analysis of α-SMA, COL-1α, and CK-14 expression at the wound site. B. Western blot analysis of protein expression in the entire wound site. C. Expression levels of IL-6 and TNF-α assessed through western blot analysis. N=4, Data are presented as means ± SD^. *^p < 0.05, ^**^p < 0.01, ^***^p < 0.001.

Additionally, Ag^+^ have been reported to modulate and suppress inflammatory responses, reducing the release of inflammatory factors such as IL-6 and TNF-α, thus mitigating wound inflammation and expediting the wound healing process[36, 39]. Consequently, we assessed the expression levels of IL-6 and TNF-α through western blot analysis. As expected, hydrogel treatment led to a significant decrease in the expression levels of IL-6 and TNF-α (**Fig 8C)**. Notably, the introduction of CLP did not result in increased expression of IL-6 and TNF-α when compared to BSA alone, indicating that CLP does not trigger additional immune responses (**Fig 8C)**.

## 3. Discussion

In our study, we introduced an innovative strategy to enhance the functionality of protein hydrogels by combining structural and biofunctional proteins without interfering with their inherent functions. BSA served as the structural protein, well-known for its role as a skeleton protein with exceptional mechanical properties. In contrast, CLP functioned as the biofunctional protein, and we crosslinked it with BSA using an Ag-S bond formed by the free cysteines of CLP. BSA underwent thiolation of lysine residues using Traut’s reagent for modification, and the thiolated protein was then crosslinked with Ag^+^ to create two types of hydrogels: BSA-Ag-BSA and BSA-Ag-CLP.

The CLP utilized in this study is a genetically engineered intrinsically disordered protein derived from the Scl-2 protein of *Streptococcus pyogenes*[1, 40, 41]. Although CLP was initially considered a “blank template” triple-helix protein, we introduced functional modifications like RGD and CPPC polypeptides, which resulted in reported biological functions[24, 42]. The substantial length of CLP’s intrinsically disordered coil sequence allowed the introduction of paired cysteines without disrupting the protein’s structure or interfering with disulfide bond formation, thus facilitating crosslinking with Ag+ through the free cysteines.

F Regarding the crosslinking strategy, Traut’s reagent-based modification has been employed in the thiolation of gelatin and proteins[43], such as ovalbumin[43], human decellularized extracellular matrix (human dECM)[29] and BSA[14, 17]. Due to the strong nucleophilic nature of thiol groups, thiolated proteins commonly undergo self-crosslinking through disulfide bonds [29] or with the assistance of Ag^+^[^14, 17]^. Some studies have explored crosslinking thiolated proteins with polypeptides to introduce additional biological functions into hydrogels[14]. However, limited research has been conducted on the crosslinking of proteins to create protein-protein hydrogels. To the best of our knowledge, there have been few reports on the development of hydrogels crosslinked by thiolated proteins and cysteine-free proteins. In our study, we genetically engineered a cysteine-free collagen-like protein (CLP) and crosslinked it with thiolated BSA using Ag^+^. Therefore, we believe that the thiolated lysine-based protein crosslinking strategy not only allows for the formation of self-crosslinked protein hydrogels but also facilitates the incorporation of functional proteins into the preparation of protein-protein hydrogels.

As for the mechanical properties of the hydrogels, BSA-Ag-CLP hydrogel exhibited comparable characteristics with BSA-Ag-BSA hydrogel. We attribute this to BSA’s role as the skeleton protein in the system, as the CLP, being an intrinsically disordered protein with a long coil sequence, couldn’t provide sufficient mechanical properties alone[44]. The rheological test showed that the inclusion of CLP resulted in a reduction in mechanical properties compared to the BSA self-crosslinked hydrogel. This reduction may be due to the flexible and intrinsically disordered structure of the triple-helical domains within CLP. The presence of CLP introduced a more flexible linker into the hydrogel system, leading to a decrease in mechanical properties. Further research is needed to investigate the relationship between protein structure and mechanical properties.

Interestingly, we observed an enhancement in biological functions, such as improved cell viability and the promotion of wound healing in both zebrafish and mouse models. Despite CLP being initially considered a “blank template” protein[1], genetic modifications and designs have improved the biological functions of the introduced CLP, as demonstrated in our previous studies[24]. The introduction of RGD sequences to improve cell adhesion and CPPC for further stabilizing the triple-helix structure of CLP’s collagen-like domain played a vital role in enhancing the biofunctional properties of CLP[42]. Therefore, CLP was utilized as a biofunctional protein in our study. This underscores the potential utility of the genetically designed CLP as a biofunctional protein in protein-protein hydrogels, enabling the incorporation of functional proteins and structural proteins into the hydrogel system. The introduction of additional biofunctional proteins or structural proteins with enhanced biological activities and clinical studies can further contribute to the development of protein-protein hydrogels.

## 4. Conclusion

In conclusion, our study advances the development of protein-protein hydrogels with enhanced functions and mechanisms. This approach not only improves the functionality of protein hydrogels but also provides a versatile method for creating protein-protein hydrogels without disrupting their inherent functions. The combination of structural control and enhanced biological functionality holds promise for diverse applications in tissue engineering and regenerative medicine. Future research should concentrate on exploring the relationship between protein structure and mechanical properties to optimize hydrogel design for specific clinical applications.

## 5. Material and methods

### 5.1 Chemical reagents

Plasmids and primers were synthesized by Sangon Biotech Co., Ltd and subsequently re-constructed in our laboratory. All the enzymes used, including ligases, endonucleases, and polymerases, were procured from Thermo Scientific. Antibodies for Collagen-1α (Catlog: Ab270993), Cytokeratin-14 (CAS: Ab119695), Anti-His (CAS: Ab18184), IL-6 (CAS: Ab 290735) and TNF-α (CAS: Ab183218) as well as the Goat Anti-Mouse IgG H&L (Alexa Fluor® 488) (CAS: Ab150113), were obtained from Abcam. α-SMA (CAS: GB111364-100) from ServiceBio Co.,Ltd. The CCK-8 and Live/Dead staining kit were supplied by Beijing Solarbio Science & Technology Co., Ltd.

### 5.2 Plasmid construction and protein purification

The cystines free collagen-like protein (CLP) were constructed as our previously reported[24]. Briefly, and pre-designed CLP genes were digested by BamHI and HindIII restriction enzymes of Thermo Scientific and colon into pET-28a (+) vector by T4 ligase.

After sequencing, the plasmid was transformed into BL21(DE3) E. coli strain. Protein expression was conducted in shake flasks using 6 L of Luria–Bertani liquid medium (LB medium) supplemented with 50 μg mL^-1^ of kanamycin at 37 °C. Once the optical density at 600 nm (OD600) reached 0.8, the cells were induced with β-D-Thio-galactoside (IPTG) at a final concentration of 0.3 mM and incubated overnight at 16 °C. After a 32-hour incubation period, the cells were harvested by centrifugation at 8000g for 15 minutes at 4 °C. After centrifugation, the cell pellet was resuspended in a lysis buffer (50 mM Tris-HCl, pH 8.0, 300 mM NaCl, 5% glycerol, v/v). The resuspended cells were sonicated to disrupt cell membranes, followed by centrifugation at 12,000 rpm for 60 minutes at 4 °C. The resulting pellet was resuspended in a binding buffer and filtered through a 0.45 μm filter to remove debris. The protein of interest was purified using a Ni^2+^ affinity column with a gradient elution of 300 mM imidazole in a buffer containing 50 mM Tris-HCl (pH 8.0) and 300 mM NaCl. The eluted protein underwent dialysis in buffer Q (20 mM Tris-HCl, pH 8.0, 50 mM NaCl, 1 mM EDTA, 1 mM Dithiothreitol (DTT), and 5% glycerol, v/v) and was loaded onto a HiTrap Capto Q column (GE Healthcare). Further purification involved a NaCl gradient (0–1000 mM) in buffer Q. Protein purity was confirmed via sodium dodecyl sulfate-polyacrylamide gel electrophoresis (SDS-PAGE) with Coomassie Brilliant Blue G-250 staining. Additionally, purified proteins were validated through Western Blot analysis with an anti-Histag antibody. Protein identification was further verified by MALDI-TOF (5800 MALDI-TOF/TOF).

### 5.3 BSA modification and hydrogels preparation

To be more specific, 0.6g of BSA (Biosharp, CAS: 9048-46-8, China) was initially dissolved in 24 mL of 0.1 M borate buffer (pH 8.0) containing 5 mM EDTA with continuous stirring on a magnetic stirrer. The amount of Traut’s reagent (Macklin, CAS: 4781-83-3, China) needed for thiolation was calculated based on the efficiency of thiolation, requiring 3.2 mL of 20% w/w Traut’s reagent for 0.6g of BSA. The Traut’s reagent was dissolved in the same solvent as BSA and was slowly added dropwise to the BSA solution. The reaction was allowed to proceed for three hours to yield a uniform BSA-SH solution, with the entire process conducted under an ice-water bath.

Following the reaction, the BSA-SH solution was concentrated using two 30 kDa ultrafiltration centrifuge tubes (Millipore UFC910096 American). Sodium acetate acetic acid buffer (1×) was added to the concentrator tube for buffer exchange, and this process was repeated three times, ultimately yielding the precursor for pure BSA hydrogel. The resulting precursor was divided into two 2 mL Eppendorf tubes, with 1 mL in each. One tube received 150 μL of 500 mM AgNO_3_ solution (Macklin, CAS: 7761-88-8, China) and was immediately shaken vigorously for 15 seconds to obtain the BSA-Ag-BSA hydrogel. The other tube received 200 μL of CLP protein and was mixed by shaking. Subsequently, 180 μL of 500 mM AgNO_3_ solution was added to this tube, and it was immediately shaken for 15 seconds to obtain the BSA-Ag-CLP mixed hydrogels.

### 5.4 Hydrogels characterizations

The hydrogel precursors (without AgNO_3_) and the hydrogels were separately loaded into syringes and injected into transparent small glass vials. This visually demonstrated the gelation of the hydrogels. Additionally, the hydrogels were injected into various molds to create different shapes, indicating their high malleability. To assess the adhesive properties of the hydrogels, they were affixed to bovine skin from which hair had been removed, and by altering the shape of the bovine skin, the adhesive properties of the hydrogel were verified. The hydrogels were dyed with various colors using dyes, pieced together, and after some time, it was observed that they had merged at the junctions. Subsequently, they were stretched to demonstrate their high extensibility. All of the aforementioned results were captured using a camera.

The hydrogel samples were freeze-dried in a vacuum freeze-dryer (Foring LGJ-10C, China) for 36 hours. Subsequently, after gold sputtering treatment using an ion sputtering apparatus (Model SC7620), they were observed under a scanning electron microscope (SEM) (Hitachi, SU8010) to examine the internal structure.

Freeze-dried hydrogels were mixed with KBr at a 1:100 ratio, followed by grinding and pressing into pellets. FTIR scans ranging from 500 to 3500 nm were conducted on these pellets using a Nicolet FTIR spectrometer (USA).

The rheological properties of the samples were tested using a rotational rheometer (MCR92, Austria). A suitable amount of the sample was placed on the sample stage, and a parallel plate rotor with a diameter of 50mm and a 1mm gap was used. For the strain modulus curve, the strain range was set from 0.01% to 1000%, and data points were collected logarithmically. The frequency was set at 1 Hz. Rheological studies were conducted in oscillatory mode with a frequency of 1 Hz, and the sample underwent multiple steps: 10% strain for 60 seconds, 800% strain for 60 seconds, 10% strain for 60 seconds, 800% strain for 60 seconds, and 10% strain for 60 seconds.

A Bio-Logic MOS-500 circular dichrois (CD) spectropolarimeter was used to examine the secondary structure of proteins. The protein samples before gelation or hydrogels were diluted to 0.1% with a 1× sodium acetate buffer, followed by CD scanning in the range of 190 to 260 nm.

### 5.5 Antibacterial properties

We evaluated the antibacterial activity of the hydrogels against gram-negative *E. coli* and gram-positive *S. aureus* using established protocols[45]. Briefly, after inoculating *E. coli* and *S. aureus*, three circular holes were created on each culture medium using a hole punch. Each hole was filled with 0.1g of hydrogel. The plates were then placed in a 37°C constant temperature incubator, and the size of the inhibition zones was observed at 12 hours and 24 hours. For OD methods, 2 ml of LB liquid culture medium was taken, and 0.1g of hydrogel was added to it. Then, 30 uL of *E. coli* and *S. aureus* were inoculated. At time intervals of 2, 4, 6, 8, 10, 20, and 24 hours, 200 uL of bacterial culture was taken to measure its OD600 value, and an equal volume of fresh LB culture medium was added. All experiments with three times repeats.

### 5.6 Cell viability, migration and immunofluorescence

To evaluate cell viability, NIH-3T3 cells and HaCat cells were cultured in DMEM (Gibco, Catalog no. 11966025) medium supplemented with 1% P/S (Gibco, Catalog no. 2441834) and 10% fetal bovine serum (Gibco, Catalog no. 10100147) at 37 °C in a 5% CO2 atmosphere. Cell viability was assessed using the CCK-8 cell counting kit. NIH-3T3 and HaCaT cells were seeded in 96-well plates and cultured until they reached an appropriate density. The cells were then cultured in culture medium containing hydrogels at concentrations of 1%, 3%, and 5% for 24 hours. Fresh culture medium containing 10% CCK-8 solution was added to each well and incubated for 30 minutes. The absorbance at 450 nm was recorded using a microplate reader (Biotek). All tests were performed with six replicates.

Further cell cytotoxicity assessment was performed by live/dead staining[24]. NIH-3T3 and HaCaT cells were seeded in 96-well plates and cultured until they reached an appropriate density over one day. Subsequently, the cells were incubated with DMEM culture medium containing 5% hydrogel. After 24 hours of culture, a Live/Dead Kit (Beijing Solarbio) was used to treat each well with 0.05% Calcein-AM (green fluorescence) and Ethidium homodimer-1 (red fluorescence) to assess cell viability. After a 5-minute incubation, fluorescence images were captured using a microscope (Leica DMi8).

For the evaluation of cell migration after hydrogel treatment, NIH-3T3 cells were cultivated in 24-well plates and subjected to scratch assays using a 1.5 mL pipette tip. Afterwards, 5% (v/v) hydrogels were introduced and co-cultured with the cells. Time-lapse images of the cells were recorded at 0, 12, 24, and 36 hours. Fiji software was employed to perform a quantitative analysis of the captured images, assessing cell migration.

Immunofluorescence was employed to assess the protein expression levels following hydrogel treatment. NIH-3T3 cells were cultured in 24-well plates under standard conditions. These cells were exposed to a 5% (v/v) hydrogel as per the experimental design. After 24 hours of incubation, the cells were fixed using a 4% paraformaldehyde solution in phosphate-buffered saline (PBS) for 15 minutes. To enhance antibody penetration, cells were permeabilized with 0.1% Triton X-100 in PBS for 10 minutes. Non-specific binding was blocked by treating cells with a 3% bovine serum albumin (BSA) solution in PBS for 1 hour. Next, the cells were incubated overnight at 4°C with primary antibodies specific to the target proteins (e.g., α-SMA, CK-14, COL-1α) at the appropriate dilution. After rinsing with PBS, cells were exposed to fluorochrome-conjugated secondary antibodies for 1 hour at room temperature. Following another round of washing to remove excess secondary antibodies, the nuclei were stained with 4’,6-diamidino-2-phenylindole (DAPI) for 10 minutes. Fluorescence images were captured using a fluorescence microscope. The obtained images were quantitatively analyzed using Fiji software.

### 5.7 Hemolysis assay and hemostasis test

Hemolysis assessments were carried out in vitro using a mouse erythrocyte suspension, following established protocols[37]. In brief, erythrocytes were obtained from whole blood through centrifugation (1000 rpm), underwent three washes with normal saline, and were then diluted to a 2% (v/v) concentration with normal saline. Subsequently, 5% (v/v) hydrogels were introduced to the erythrocyte suspensions (1 mL) and incubated at 37 °C for 1 hour. As negative and positive controls, normal saline and Triton X-100 were used in place of the hydrogels, respectively. After the incubation, the erythrocyte suspensions were centrifuged at 1000 rpm for 15 minutes. The supernatant was collected, and the absorbance was measured at 545 nm using a UV–vis spectrophotometer.

The hemostatic potential of the hydrogels was evaluated using a hemorrhaging liver model (BALB/c mice, 25–30 g, 10–12 weeks old, male)[37]. In brief, mice were anesthetized with isoflurane administered via a respiratory anesthesia system. The mouse’s liver was exposed through an abdominal incision, and any residual fluid on the liver’s surface was gently removed. A pre-weighed filter paper was positioned beneath the liver. Following the induction of liver hemorrhaging with a 21 G needle, 50 μL of each hydrogel was immediately injected onto the bleeding site. A control group underwent no treatment after the liver was punctured, and gauze was applied. Images were captured after 5 minutes of bleeding and subsequently analyzed using Fiji software.

### 5.8 The wound healing of zebrafish

Adult zebrafish of the wild-type AB strain, aged approximately 3 to 4 months, were raised in a recirculating aquatic system at a controlled temperature of 28.5 °C, following a 10/14-hour dark/light cycle, in accordance with established standards. After an acclimatization period of one week, the zebrafish were categorized into three groups, each consisting of six individuals: the control group, the BSA-Ag-BSA group, and the BSA-Ag-CLP group. For the wound healing experiment, each zebrafish was anesthetized individually by immersing them in a 0.2% MS-222 solution. Once anesthetized, a full-thickness wound with an approximate diameter of 2 mm was generated on the left flank of the zebrafish, just anterior to the anal and dorsal fins, using a UV-laser beam set at 3 watts for 5 seconds. A 0.1% methylene blue solution was applied to the wound, and images were captured with a stereomicroscope. Subsequently, the appropriate experimental adhesive sheet was applied to the wound. Following this procedure, the zebrafish were allowed to recover for 5 minutes in the system water before being transferred to their respective experimental tanks. The experimental solution in these tanks was renewed every 2 days. At specific time points, specifically at 0, 3, 7, 13, and 18 days after wounding, the fish were temporarily removed from their test tanks and allowed to recover in system water for 5 minutes. Subsequently, the fish were anesthetized, and images of the wounds were captured using a microscope. The wound area was carefully monitored and measured at these predetermined time intervals. On the 18th day, zebrafish from each group were euthanized, and the wound areas were collected and placed into 4% paraformaldehyde (PFA) (Solarbio P1110, China) for 24 hours to fix the tissues. After embedding in paraffin, 5 μm thick sections were prepared, followed by H&E (Hematoxylin and Eosin) and Masson staining.

### 5.9 Mouse diabetic wound model

Male Balb/c mice weighing 20-25g were sourced from Charles River and acclimatized to the laboratory environment for 7 days before the commencement of the experiment. Diabetes was induced through a one-time intraperitoneal injection of 110 mg kg^−1^ streptozotocin (Step, Sigma Aldrich) dissolved in citrate buffer (pH 4.5) over a single day[35]. After 7 days of Step treatment, blood glucose levels in the mice were measured. Mice with consistent glucose levels exceeding 10.0 mmol L^−1^ were considered to be diabetic.

Once the diabetic mouse model was established, mice were anesthetized using isoflurane (RWD, R510-22) administered via a respiratory anesthesia system (RWD, R510IP). The mice were positioned in a prone stance, their fur was shaved, and the back skin was sterilized with iodine tincture and cleaned with 75% ethanol. A circular wound with a 1 cm diameter was created on the back skin of each mouse. Subsequently, BSA-Ag-BSA and BSA-Ag-CLP hydrogels were applied to the wounds. Imaging was conducted on days 0, 3, 7, and 12 following treatments. The wound areas were analyzed using Fiji software, and the wound healing rate was calculated using the formula: Wound healing rate (%) = (S0 - Sn) / S0 × 100, where S0 and Sn represent the wound area on day 0 and day n after treatment. All animal experiments were performed in compliance with the guidelines and regulations approved by the Inner Mongolia University Ethics Committee.

### 5.10 Histology and immunohistochemistry

After 12 days, the mice were euthanized with CO2, and their wound skin samples were collected. These tissue samples were fixed in 4% paraformaldehyde for 24 hours and processed using standard haematoxylin-eosin (H&E) staining and Masson’s trichrome staining. We observed the stained sections under an optical microscope (Leica, DMi8 S). For immunohistochemistry, we followed established methods[24, 46]. Antigen retrieval was performed at 95 °C for 1 hour. The sections were then incubated with primary antibodies targeting collagen-1α and cytokeratin-14 at 4 °C. After that, a secondary antibody (anti-rabbit/anti-mouse Envision) was applied for 30 minutes. Finally, we used fluorescence microscopy (Leica, DMi8 S) to view and analyze the immunostained sections.

### 5.11 Western bolt

The wound site was lysed to extract total proteins, and their concentrations were determined using a BCA protein assay kit. Subsequently, 30 μg of protein samples were separated on 10% SDS-PAGE gels and transferred onto nitrocellulose membranes. Primary antibodies against COL-1α, CK-14, α-SMA, IL-6 and TNF-α (dilution 1:500, Abcam) were incubated with the membranes overnight at 4 °C, with β-actin used as the internal reference. The membranes were later exposed to a secondary antibody, anti-rabbit IgG horseradish peroxidase-conjugated (dilution 1:5000). Protein bands were visualized using an Enhanced Pico Light Chemiluminescence Kit, and densitometric analysis was conducted with Image J software.

### 5.12 Statistical analysis

In this study, the data is expressed as mean ± standard deviation. Statistical analysis was carried out with GraphPad Prism 8 software, employing one-way analysis of variance (ANOVA). A *p*-value below 0.05 was considered statistically significant, denoting a notable difference between the groups. Image analysis was executed using Fiji software, ensuring precise and dependable measurements and quantification of the images.

## 6. Acknowledgement

This project was supported by a grant from National Natural Science Foundation (22365022); Inner Mongolia Natural Science Foundation (2022QN03014); Inner Mongolia Science and Technology Project (2022ZY0050); Inner Mongolia Youth Science and Technology Talent Support Project (NUYT23092); Science and Technology Leading Talent Team in Inner Mongolia Autonomous Region (2022LJRC0009); Inner Mongolia Autonomous Region Science and Technology Project (2023YFHH0010). X.L, J.W and Z.G contributed equally to this work. All authors have approved the final version of this manuscript. Ethical approval for the in vivo diabetic wound healing experiments was granted by the Institutional Animal Care and Use Committee of Inner Mongolia University.

## 8. Data Availability Statement

The data that support the findings of this study are available from the corresponding author upon reasonable request.

