## Supplementary material for "Thiolation-Based Protein-Protein Hydrogels for Improved Wound Healing": supplementary information.pdf

MCGGADEQEEKAKVRTELIQELAQGLGGIEKKNFPTLGDEDLDHTYMTKLLTYLQEREQ

AENSWRKRLLKGIQDHALDGGPCPPCGPKGEQGPQGLPGKDGEAGAQGPAGPMGPAGE

QGEKGEPGTQGAKEDRGETGPKGPKGERGEAGPAGKDGEPGPVGPAGPKGEQGPQGLPG

KDGEAGAQGPAGPMGPAGEQGEKGEPGTQGAKEDRGETGPKGPKGERGEAGPAGKDGE

PGPVGPAGPKGEQGPQGLPGKDGEAGAQGPAGPMGPAGEQGEKGEPGTQGAKEDRGET

GPKGPKGERGEAGPAGKDGEPGPVGPAGGPCPPCRGDGGC*

**Supplementary Figure 1:** Amino acid sequence of the genetically encoded CLP. CGG; V-domain; CPPC; Collagen-like domain-1; Collagen-like domain-2; Collagen-like domain-3; RGD.

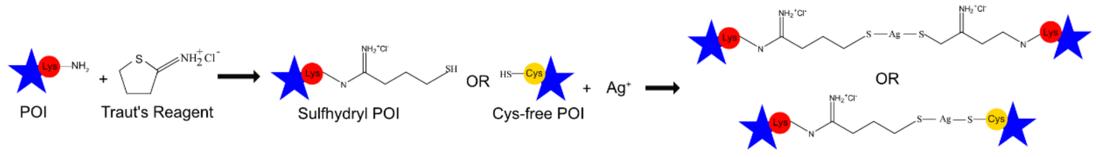


**Supplementary Figure 2:** Traut's reagent-based lysine modification and mechanisms of hydrogel formation.

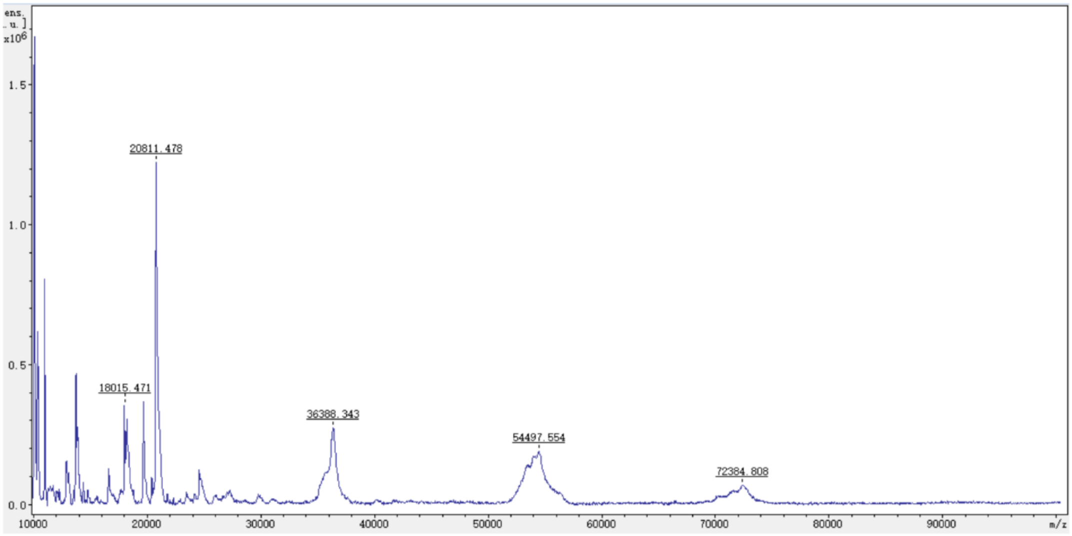


**Supplementary Figure 3: Maldi-TOF of the recombined CLP.**

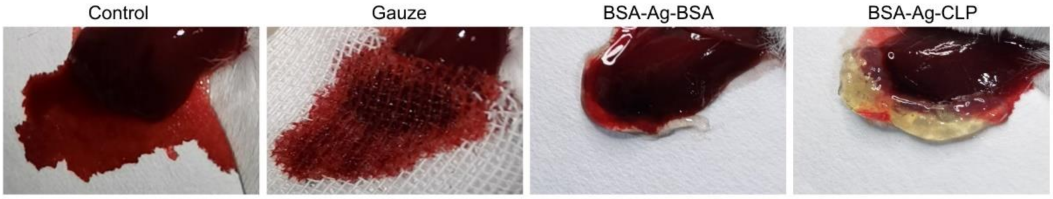


**Supplementary Figure 4: Images of applying hydrogels for in vivo hemostasis (n = 3).**
